# Layer 5 Martinotti and pyramidal neurons encode spatial information in the primary motor cortex

**DOI:** 10.64898/2026.02.25.707928

**Authors:** Barbara Ciralli, Thawann Malfatti, Klas Kullander

**Affiliations:** Department of Immunology, genetics and pathology, Uppsala University, Uppsala, Sweden

**Keywords:** Martinotti alpha 2 cell, pyramidal cell, Spatial information, Place cell, Grid cell, Head-direction cell

## Abstract

Mammals rely on external sensory cues and internal bodily signals to create a precise spatial awareness. The motor cortex is essential for this process, integrating proprioceptive and spatial information. Here, we investigate how layer 5 martinotti (M*α*2) interneurons and pyramidal cells contribute to spatial processing in primary motor cortex (M1). Using miniscopes in freely moving mice expressing CaMKIIa-GCaMP8f or DIO-GCaMP8f in Chrna2-Cre+ mice, we recorded activity from both cell types. We found that both M*α*2 and pyramidal cells are tuned to place, head direction, and grid fields, with distinct characteristics. Notably, both cell types exhibit an overrepresentation of co-tuned place- and head-direction responses compared to chance. Moverover, pyramidal head direction-tuned cells show sharper tuning than M*α*2 head direction-tuned cells. Further, place-tuned Mα2 cells are more active, whereas pyramidal place-tuned cells showed reduced activity compared to non-place-tuned ones. Additionally, both cell types exhibit lower activity in head-direction-tuned compared to non-head direction-tuned cells. These findings suggest that distinct neural coding strategies are employed by different cell types within the M1 for processing spatial information, highlighting the complex neural mechanisms underlying spatial cognition.

## Introduction

As mammals navigate their surroundings, they integrate external sensory inputs with internal self-motion cues to build an accurate mental map of their environment [1]. The motor cortex contributes to this process by integrating proprioceptive and processing spatial information [2, 3]. For example, neurons in the primary motor cortex (M1) exhibit space and orientation tuning, which has been observed in recordings from monkeys navigating a room to reach a food reward. These recordings showed that M1 neurons were tuned not only to the animal’s position within the room but also to its orientation relative to the room [4]. Moreover, during locomotion, it has been demonstrated that the mouse motor cortex shows increased activity in neurons from layer 5 [5]. Using intracellular recordings, they found a greater percentage of activated layer 5b neurons (53%) compared to those that were suppressed (38%) pointing out the motor cortex participation, specifically the layer 5b during locomotion [5].

The motor cortex’s ability to adapt motor plans based on changing spatial information is essential for effective navigation and contributes significantly to our overall spatial awareness and orientation. The cortico-hippocampal network reveals that three critical regions, the retrosplenial cortex, posterior parietal cortex and secondary motor cortex (M2), serve as key hubs for integrating spatial information with motor planning [6]. These regions are intricately connected to other structures within the hippocampal formation [7, 8], forming a complex network that underlies spatial cognition. Both retrosplenial cortex and posterior parietal cortex send projections to the M2 [9], which in turn projects to M1 [10, 11], establishing a robust anatomical substrate for sensorimotor integration. Moreover, Martinotti cells in the M1 motor cortex receive direct input from thalamic nuclei, which in turn relay information from other cortical structures as well as basal ganglia and the cerebellum [11]. This hierarchical organization demonstrates the role of the M1 in integrating spatial information and translating it into motor output, ultimately enabling locomotion and exploration.

Interestingly, studies using calcium imaging have identified patterns in certain areas of the cerebral cortex, which are similar to the population code for spatial processing observed in the hippocampus [12, 13]. For instance, spatial activity has been registered in the retrosplenial cortex, which was hindered by bilateral damage to the hippocampus [13]. Furthermore, place-tuned cells and head direction-tuned cells have been described in the visual cortex [14, 15, 16, 17], posterior parietal cortex [18] and somatosensory cortex [19]. However, cells exhibiting place, grid, and head direction-like characteristics have yet to be identified in the M1, which is an important region for planning, controlling, and executing voluntary movements [20, 21, 22].

Inhibitory interneurons can encode spatial information. For example, hippocampal interneurons from CA1, dentate gyrus, and subiculum exhibit place-tuned firing fields that are similar to those of excitatory cells [23]. However, previous studies have also suggested that CA1 interneurons may have broader tuning profiles than those of the excitatory neurons they directly connect [24]. Notably, somatostatin-expressing (SOM) interneurons in the cortex are of particular interest, as their inactivation has been shown to impair spatial selectivity in place-tuned cells within the medial entorhinal cortex [25]. A subset of these SOM-positive neurons in the cortex, known as martinotti cells, have been shown to play a critical role in synchronizing cortical pyramidal cell activity [26]. These cells have gained increasing attention due to their key role in modulating cortical circuitry and influencing various aspects of neural processing. With a characteristic axonal projection that ascends toward the cortical surface, the martinotti cells provide inhibitory feedback to pyramidal neurons, thereby contributing to the regulation of excitatory transmission and the maintenance of cortical oscillations [26, 27]. Furthermore, layer 5 martinotti (M*α*2) cells receive direct input from M2 and thalamus [11]. In this study, we aimed to determine whether M*α*2 cells and layer 5 pyramidal neurons encode information related to location, head direction, and spatial grid patterns.

## Materials and methods

### Animals

All animal procedures were approved by the local animal research ethical committee (Ethical permit number: 5.8.18-07526/2023). Mice were housed with littermates in ventilated cages (up to 5 mice/cage) with bedding and enrichment (a carton house and shredded paper strips), kept in a 12-h light on/light off cycle (6 a.m.–6 p.m.), and maintained at 21 ± 2 °C with a humidity of 45-64%. Mice were provided food (diet pellets, Scanbur, Sweden) and tap water ad libitum. 8 females Chrna2-Cre tg/wt were used for imaging the M*α*2 cells [26] and 6 females Chrna2-Cre wt/wt were used for imaging the pyramidal cells in the M1 (as described below in the “Viral injections” section). Mice were 8-10 weeks old at the start of experiments. After lens implant surgery, the animals were housed individually to decrease risk of implant detachment.

### Viral injections

Mice were sedated with 1-4% Isoflurane (Baxter). Analgesia was given subcutaneously; 2 mg/kg bupivacaine (Marcain, AstraZeneca), 5 mg/kg carprofen (Norocarp vet, N-vet or Rimadyl Biovis vet) and 0.1 mg/kg buprenorphine (Vetergesic vet, Ceva). A midline incision was made in the scalp, muscles and periosteum were removed using 3% hydrogen peroxide (Sigma-Aldrich) and a hole (1 mm in diameter) was drilled at the M1 area [coordinates relative to bregma: AP 0.8 mm, ML 1.5 mm and DV -1.3 mm; 28]. Next, the virus AAV9-syn-FLEX-jGCaMP8f, for martinotti cells imaging, or AAV9-CaMKIIa-GCaMP8f, for pyramidal cells imaging, was injected using a 10 µl Nanofil Hamilton syringe (WPI, USA) mounted on a stereotaxic frame. Ten minutes post injection the needle was slowly withdrawn from the brain and the wound was stitched with resorbable sutures (Vicryl rapid, Ethicon, 6-0). Mice were then left to recover for 2 weeks before any behavioral experiments were initiated. Viral transduction was confirmed by post-hoc histological analysis of brain sections.

### Open-field recordings

The animals were recorded in an open field arena with 50 cm of diameter. A webcam (Logitech QuickCam Pro 9000) was placed in the top center of the arena, in order to record individual mice behaving freely during locomotion. The camera was recorded in sync with the miniscope through the miniscope data acquisition software.

### Lens implantation

The procedure was initially identical to the viral injections, but instead of injecting a virus, a prism lens (CLH type glass, lens diameter 1.0 mm, prism size 1.0 mm, total length of lens+prism 3.68±0.156 mm, working distance 0.3 mm, enhanced aluminum and SiO2 protective coating, Go!Foton, USA) for recordings of M1 cortical layer 5 was implanted unilaterally using the stereotaxic frame. To implant the prism lenses, a round cranial window with 1.1mm diameter was opened, with center at the coordinates 0.8 AP and 2.3 ML. Next, the tip of a dissection knife was inserted into the cortex for three minutes before the lens was lowered into the same place with the prism edge at 0.3 mm lateral and -0.5 mm dorsoventral to the viral injection coordinates (final coordinates for the edge of the lens: AP 0.8 mm, ML 1.8 mm and DV -1.8 mm). The lens was fixed to the skull and anchor screws with dental cement (light-cured flowable composite, Tetric EvoFlow). Silicon glue (Kwik-Sil, World Precision Instruments) was added to protect the lens from mechanical damage in between experiments. Mice were treated with Enrofloxacin 5 mg/kg (Baytril) for two days pre-surgery and seven days post-surgery. Two to three weeks post implantation the mice were again anesthetized, the scalp incised, and the baseplate (used to attach the miniscope during recordings) was mounted on top of the skull with cyanoacrylate glue. Lens position was confirmed by post-hoc histological analysis of brain sections.

### In-vivo calcium imaging analysis

To extract cell activity for analysis, we employed calcium imaging techniques using a viral vector with DIO- or CamKII promoter-driven expression of GCaMP8f for martinotti and pyramidal cells, respectively. As previously established [29, 30, 31], increased calcium levels correlate with enhanced cell activity, however, direct estimation of single spikes in 1-photon recordings is not feasible. We processed Miniscope videos using a standardized approach [32] as previously described [27]. Specifically, we applied a factor of 3 downsampling to reduce the video resolution while retaining essential features, and corrected for motion artifacts. Next, we extracted cells’ signals using constrained nonnegative matrix factorization for 1-photon data [CNMF-E; 33] from the downscaled and motion-corrected data. Fluorescence traces were then detrended, deconvolved, and normalized by its noise standard deviation, resulting in fluorescence traces here expressed as ΔF/F, to ensure that all signals have comparable amplitudes.

### Place, grid and head direction cells classification

#### Activity and correlation maps

First, we binned the 2D location data for each cell into a grid with a spatial bin size of 2.5 cm, resulting in a 20×20 bin map for the 50 cm open-field arena. To filter out activity unrelated to locomotor behavior, only frames where the animal’s instantaneous speed was above 2.5 cm/s were included. We then computed the occupancy by counting the amount of time the animals spent on each spatial bin and calculated the cells’ spatial activity for each visited spatial bin as the sum of its activity in that bin. The resulting spatial activity map was normalized by dividing it by the occupancy map. Unvisited spatial bins were filled with the average activity of the map, and we applied a Gaussian smoothing kernel with a standard deviation of 3 cm. Finally, to calculate the spatial autocorrelogram, we normalized each cell’s activity map to have zero mean and unit standard deviation, then computed the 2D spatial autocorrelation, representing the similarity between adjacent spatial bins in the activity map.

To analyze the spatial activity fields of place-tuned cells, we first determined a global threshold for activity within each spatial bin as the 75th percentile of the sorted activity values across all maps, normalized by the maximum activity in each animal’s maps. Then, for each cell’s activity map, we scaled the global threshold by the maximum activity value within that cell’s map, and used this local threshold to determine the locations where the cell was active. We next iterated over the spatial bins to form contiguous activity fields. For each identified spatial bin that satisfied the local threshold, we checked whether it had already been included in a previously identified field. If not, we initialized a new field and expanded it by including neighboring spatial bins that also met the threshold within a 3×3 neighborhood. This process continued recursively until all neighboring bins were processed. Once each activity field was fully expanded, the field size was quantified as the number of spatial bins multiplied by the square of the spatial bin size in centimeters. Only fields with a minimum size of 7.5 cm2 were kept. The center of each activity field was computed as the centroid of the polygon formed by the center of the spatial bins within the field.

#### Spatial Information Score

To identify place cells, we calculated the spatial information score (SIS) for each neuron based on their spatial activity map, which measures the amount of information about the animal’s position that is carried by the activity of a single neuron, as described by Skaggs et al. [34]. A high SIS indicates that the neuron has strong place field properties, meaning it fires more when the animal is in specific locations within the environment. The SIS for each neuron was calculated by summing the product of the occupation probability, the response strength of the neuron, and the natural logarithm of the response strength of the neuron for each cell in the activity map. The formula to compute the SIS is:

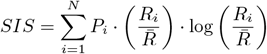

where *P*_*i*_ is the occupation probability of location bin *i, R*_*i*_ is the response amplitude of the cell at location *i*, and 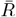 is the average of the cell activity over all visited bins.

#### Gridness

To identify grid cells, we calculated the gridness score for each neuron, which measures how well their activity can be described by a grid structure, based on a previous implementation [14]. A high score indicates that the neuron fires more when the animal is in specific locations within the environment, meaning it has strong grid field properties. The central peak of each gaussian-smoothed autocorrelogram was identified and excluded from further analysis. We then extracted radial samples from the autocorrelograms by iterating over expanding circular rings around the central peak, calculating the Pearson correlation coefficient of these samples against their rotations at specified angles categorized into Group 1 (60° and 120°) and Group 2 (30°, 90°, and 150°). The gridness score was defined as the highest minimum difference between the correlations of Group 1 and Group 2 across all circular sample radii.

#### Mean Vector Length

To identify head direction cells, we calculated the mean vector length (MVL) for each neuron based on the mouse head direction in relation to the arena, based on a previous implementation [14]. A high MVL indicates that the neuron fires more when the animal has a specific head direction within the environment, meaning it has strong head direction properties. The mean vector length was calculated for each neuron by normalizing the tuning curve, creating an angular array, and computing the X and Y components by multiplying the tuning curve by the cosine and sine of the angular array, respectively. Then, it calculates the Euclidean norm of the tuning curve and divides it by the sum of the response strengths of all angles to obtain the mean vector length. The formula to compute the MVL is:

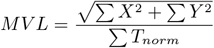

where *X* and *Y* are the components of the tuning curve, and *T*_*norm*_ is the mean response strength of all angles.

#### Chance levels

To estimate the chance level for spatial information score, gridness, and mean vector length, allowing classification of neurons as place, grid, or head direction cells, respectively, we employed a shuffling-based approach previously described [35, 14]. Specifically, each neural activity trace was time-shifted randomly within 5% to 95% of the original signal’s duration. This resulted in a new time series without spatial structure while preserving relationships between adjacent data points. A set of 200 randomly time-shifted traces was generated for each cell, and SIS, gridness, and MVL were extracted for each trace, as described for real signals. The estimated chance levels for these features were then obtained by calculating the 95th percentile values across all iterations of signal rolling from all cells within the same animal, which enables animal-specific estimation of chance levels.

### Statistical analysis

The independence of spatial feature representations was assessed using contingency table analysis. Specifically, we performed Fisher exact tests on 2×2 tables constructed from the presence or absence of place-tuned, grid-tuned, and head direction-tuned cells in each trial. We computed the observed frequencies of joint occurrences for each pair of cell types by iterating over all trials and incrementing the corresponding table entries accordingly. The Fisher exact odds ratio statistic was then calculated, which provides a measure of the likelihood that the observed frequencies arise by chance under the assumption of independence between the two variables. We performed this analysis for both M*α*2 and pyramidal cells, and for all three pairs of cell tuning: place-tuned vs grid-tuned, grid-tuned vs head direction-tuned, and place-tuned vs head direction-tuned. The effect of neural population (Ma2 vs pyramidal cell) on place cell size, grid size and grid angle was calculated using the Kruskal-Wallis test, due to their non-parametric distribution (verified using Shapiro-Wilk test). Pairwise comparisons were performed using the Mann-Whitney U test. For all violin plots, the filled area represents the data range, and horizontal lines and triangle markers represent medians and means, respectively. For all bar plots, bars represent the mean and error bars represent the standard error of the mean.

### Software availability

Calculations were done using Scipy [36], Numpy [37] and SciScripts [38], all plots were produced using Matplotlib [39] and schematics were drawn using Inkscape [40]. Cell fluorescence signals were extracted and processed from miniscope videos using CaImAn [32] and OpenCV [41]. Tracking of the mouse position in the open-field arena was done using DeepLabCut [42]. All scripts used for data analysis, including the DeepLabCut trained network, are available online [git repository; 43].

## Results

### M*α*2 cells and pyramidal neurons exhibit tuning for spatial features

To investigate the spatial information coding properties of M1 M*α*2 cells and pyramidal neurons, targeted genetically using the combination of the Chrna2-Cre+ line and a Cre-dependent GCaMP8f vector [26] and a CaMKIIa-dependent GCaMP8f vector, respectively, mice explored an open field arena while locomotion and calcium imaging data were recorded (Figure 1A-E). We determined whether these cell populations can be tuned to encode place, grid, or head direction by calculating their spatial information score (SIS), gridness and mean vector length (MVL), respectively. Interestingly, we found that both M*α*2 cells (Figure 1F, 2A-C) and pyramidal neurons (Figure 2D-F) exhibit tuning for place, grid, and head-direction. Given that the chance levels of spatial feature coding are animal-specific, we evaluated the distribution of scores for tuned and untuned cells for each feature to ensure that tuned populations indeed have larger scores than untuned populations upon pulling cells from multiple animals. Indeed, our results showed that scores were significantly higher in tuned cells compared to untuned cells for each feature in both M*α*2 (Mann-Whitney U eff. size > 0.337, p < 3.1e-31; Figure 2A-C top) and pyramidal cells (eff. size > 0.352, p < 1.1e-16; Figure 2D-F top), with tuned cells showing histogram distributions above chance distribution (shuffles, Figure 2A-F bottom).

**Figure 1:**
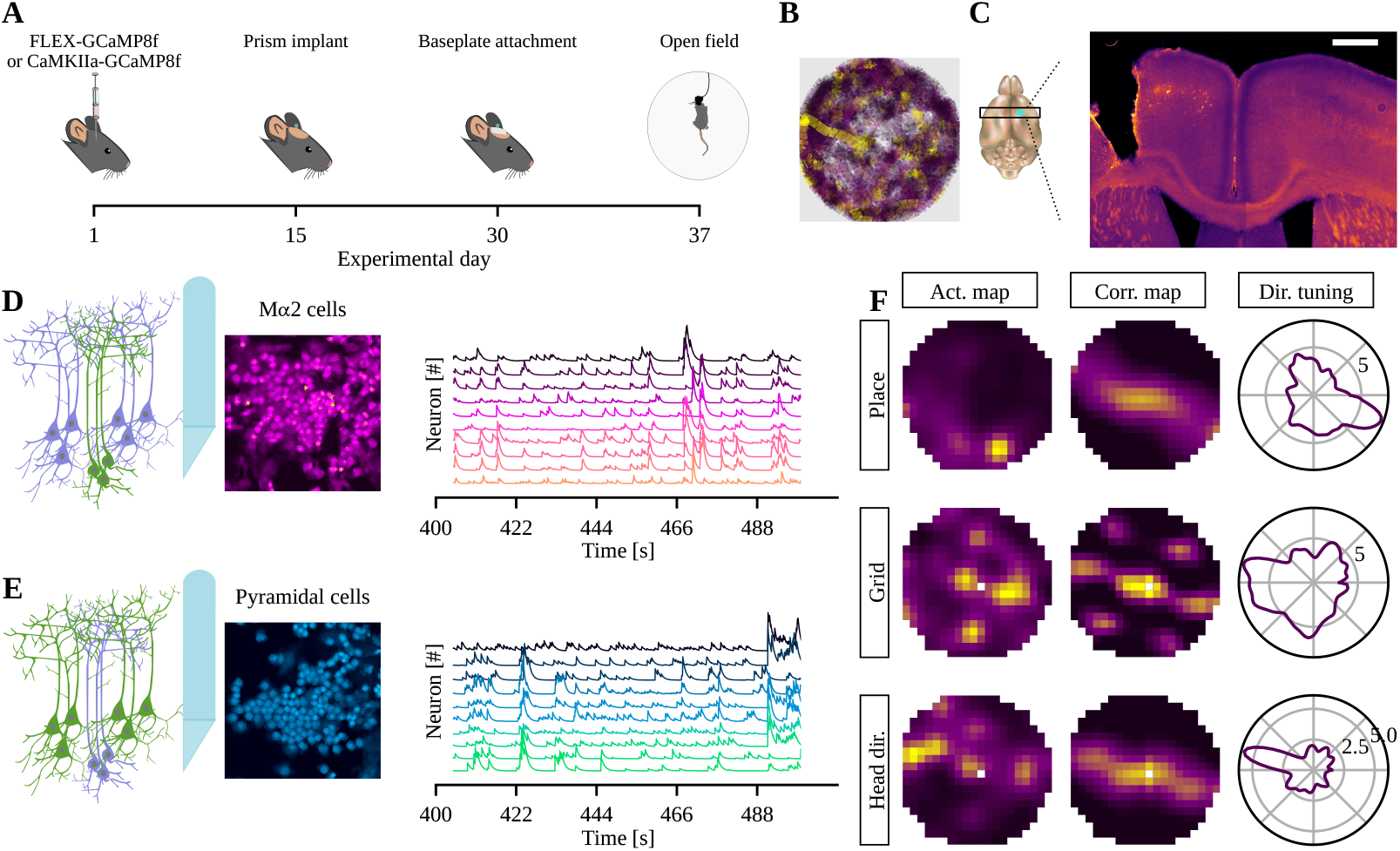
Experimental timeline and representative examples. A) Experimental timeline depicting the time interval between each surgery and the open-field recording. B) Example of tracking of one animal in the open-field arena, with colormap representing the normalized average activity of a grid-tuned neuron. C) Schematic of lens placement in the brain (left) and a representative brain section showing Ma2 expression in the M1 layer 5 and the prism lens positioning. Scalebar: 500µm. D-E) Schematics representing the recorded cells (left), representative spatial footprints (middle) and normalized calcium fluorescence traces (right) for M*α*2 (D) and pyramidal cells (E). F) Activity map (left), spatial correlation map (center) and head direction tuning (right) of place cell (top), grid cell (center) and head-direction cell (bottom).

**Figure 2:**
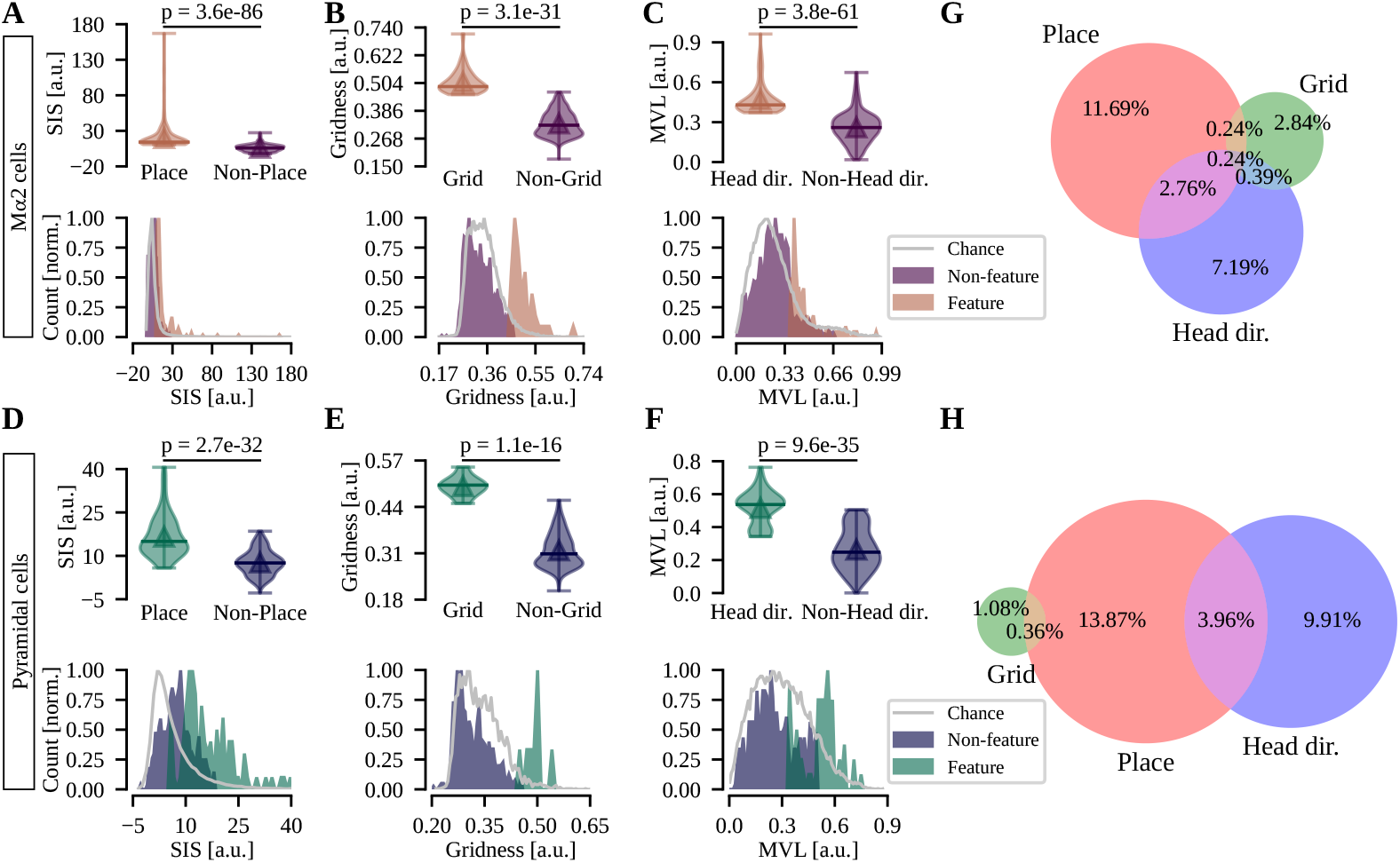
Both M1 M*α*2 and pyramidal cells can encode place, grid and head direction information. A-F) Top, spatial information score (SIS), gridness and mean vector length (MVL), separated by place-tuned, grid-tuned and head direction-tuned cells, respectively for M*α*2 cells (A-C) and pyramidal cells (D-F). Bottom, Normalized histograms for each feature showing the distributions for shuffles (chance, silver), non-feature and feature cells. G-H) Venn diagrams showing the proportion and overlap of M*α*2 (G) and pyramidal (H) cells encoding for spatial features.

Further analysis revealed that a subset of spatially-tuned M*α*2 cells were more likely to be tuned for place than grid or head direction. Specifically, an average of 14.93% of recorded cells (n=8 mice, 189/1266 cells) exhibited tuning for place, whereas 3.71% (47/1266 cells) and 10.58% (134/1266 cells) showed tuning for grid and head direction, respectively (Figure 2G). Similarly, pyramidal neurons were also more likely to be tuned for place than grid or head direction. On average, 18.2% of recorded cells (n=6 mice, 101/555 cells) exhibited tuning for place, whereas 1.44% (8/555 cells) and 14.05% (78/555 cells) showed tuning for grid and head direction, respectively (Figure 2H).

### Place and head-direction tuning are not independent in both M*α*2 cells and pyramidal cells

Due to overlaps observed in the feature diagrams of both M*α*2 and pyramidal cells, we next evaluated whether the coding of multiple spatial features is independent from one another. This analysis revealed that the observed co-occurrences between pairs of spatial features deviated significantly from those expected under independence for M*α*2 cells (Fisher exact odds ratio = 2.572, p = 1.8e-05; Table 1, left). Specifically, we found that the proportion of cells that are neither place-tuned nor head direction-tuned was slightly elevated (77.49% vs 76.07%), whereas the proportions of place-tuned-only and head direction-tuned-only were reduced (11.93% and 7.58%, compared to 13.35% and 9.0%, respectively). Conversely, the proportion of cells that are classified both as place-tuned and head direction-tuned was higher than expected (3.0% vs 1.58%).

**Table 1.** Contingency table for M*α*2 and pyramidal cells spatial features distribution.

| Place x HD |  |  |  |  |
| --- | --- | --- | --- | --- |
|  | Mα2 cells (observed / expected) |  | Pyramidal cells (observed / expected) |  |
|  | Not HD | HD | Not HD | HD |
| Not Place | 77.49 / 76.07 | 7.58 / 9.0 | 71.89 / 70.45 | 9.91 / 11.35 |
| Place | 11.93 / 13.35 | 3.0 / 1.58 | 14.23 / 15.67 | 3.96 / 2.52 |
| Fisher exact odds ratio | 2.572; p = 1.8e-05 *** |  | 2.02; p = 1.6e-02 * |  |

| Place x Grid |  |  |  |  |
| --- | --- | --- | --- | --- |
|  | Mα2 cells (observed / expected) |  | Pyramidal cells (observed / expected) |  |
|  | Not Grid | Grid | Not Grid | Grid |
| Not Place | 81.83 / 81.91 | 3.24 / 3.16 | 80.72 / 80.62 | 1.08 / 1.18 |
| Place | 14.45 / 14.37 | 0.47 / 0.55 | 17.84 / 17.94 | 0.36 / 0.26 |
| Fisher exact odds ratio | 0.828; p = 0.835 |  | 1.508; p = 0.642 |  |

| Grid x HD |  |  |  |  |
| --- | --- | --- | --- | --- |
|  | Mα2 cells (observed / expected) |  | Pyramidal cells (observed / expected) |  |
|  | Not HD | HD | Not HD | HD |
| Not Grid | 86.33 / 86.1 | 9.95 / 10.19 | 84.68 / 84.88 | 13.87 / 13.67 |
| Grid | 3.08 / 3.32 | 0.63 / 0.39 | 1.44 / 1.24 | 0.0 / 0.2 |
| Fisher exact odds ratio | 1.779; p = 0.147 |  | 0.0; p = 0.607 |  |

Similarly, from all pairwise combinations of place-tuned, grid-tuned and head direction-tuned cells, we found frequencies of place-tuned vs head direction-tuned different from expected by chance for pyramidal cells (Fisher exact odds ratio = 2.02, p = 1.6e-02; Table 1, right). The contingency table shows a slight increase in cells that are neither place-tuned nor head direction-tuned (71.89% vs 70.45%), while the percentage of place-tuned-only and head direction-tuned-only were reduced (14.23% and 9.91%, compared to 15.67% and 11.35%, respectively). In contrast, cells classified both as place-tuned and head direction-tuned were more present than expected by chance (3.96% vs 2.52%).

Interestingly, this was the only pair of spatial features to show significant contingency, as we found no deviations from independence between place-tuned and grid-tuned, or between grid-tuned and head direction-tuned cells, for both M*α*2 cells and pyramidal cells. This suggests that the tuning of some cells to place is not independent of their tuning to head direction, suggesting a more integrated representation of spatial information in M1 neurons.

Further, we extracted one measure for place-tuned cells, the size of the activity place field, and two measures for grid-tuned cells: the size (defined as the average distance of the first neighbor fields from the center field in the autocorrelation map) and the phase angle (defined as the average angle formed between the first neighbors and the center of the autocorrelation map) of the grid. We found no difference in place field sizes between M*α*2 and pyramidal cells (157.716 ± 10.744 cm^2^ vs 136.815 ± 11.892 cm^2^; Kruskal-Wallis, F(1,216) = 1.234, p = 0.268; Figure 3A-C), showing that M*α*2 and pyramidal place-tuned cells are activated by similar spatial areas. Similarly, the size of the grid in grid-tuned cells was comparable between M*α*2 and pyramidal cells (25.393 ± 1.504 cm vs 21.347 ± 1.546 cm; F(1,41) = 2.145, p = 0.151; Figure 3D), and the grid orientation was also similar (29.666 ± 2.792º vs 34.723 ± 7.761º; F(1,41) = 0.552, p = 0.462; Figure 3E). These findings suggest that both M*α*2 and pyramidal cells in the motor cortex share similar place field size and grid parameters, supporting the idea that distinct cell types may still operate within a common spatial processing mechanism.

**Figure 3:**
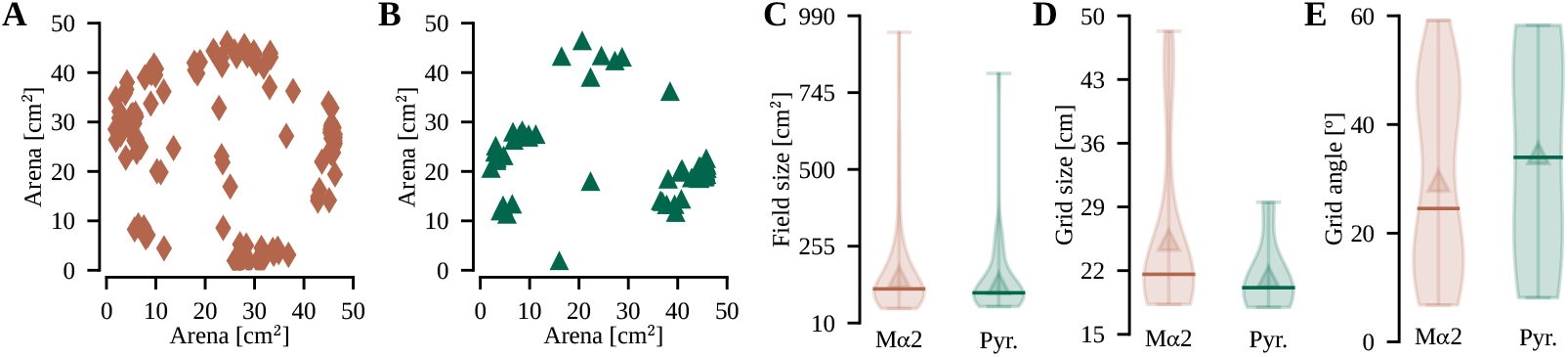
Place and grid features are similar between pyramidal and M*α*2 tuned cells. A-B) Center of activity fields for place-tuned M*α*2 (A) and pyramidal cells (B). C) Activity field size for M*α*2 (orange) and pyramidal cells (green) place-tuned cells. D-E) Grid size and phase angle for M*α*2 (orange) and pyramidal cells (green) grid-tuned cells.

### Pyramidal cells encode more head direction information than M*α*2 cells

We next evaluated the tuning of M*α*2 cells and pyramidal neurons by comparing their spatial feature scores in more detail. Although we found that M*α*2 cells have a lower mean percentage of place-tuned and head direction-tuned cells, but a higher mean percentage of grid-tuned cells compared to pyramidal cells, no differences were found in the overall percentage of tuned cells between the two populations (n = 8 and 6 mice respectively; Figure 4A-C). However, when investigating whether there are differences between the tuning of M*α*2 cells and pyramidal neurons from all animals (Figure 4D-F), we found that M*α*2 head direction-tuned cells display lower MVL compared to pyramidal head direction-tuned cells (Mann-Whitney U eff. size = 0.252, p < 2.5e-04; Figure 4F). No difference in tuning was found between M*α*2 and pyramidal place-tuned and grid-tuned cells (Figure 4D-E). This suggests that M*α*2 cell activity encodes less head-direction information than pyramidal cells.

**Figure 4:**
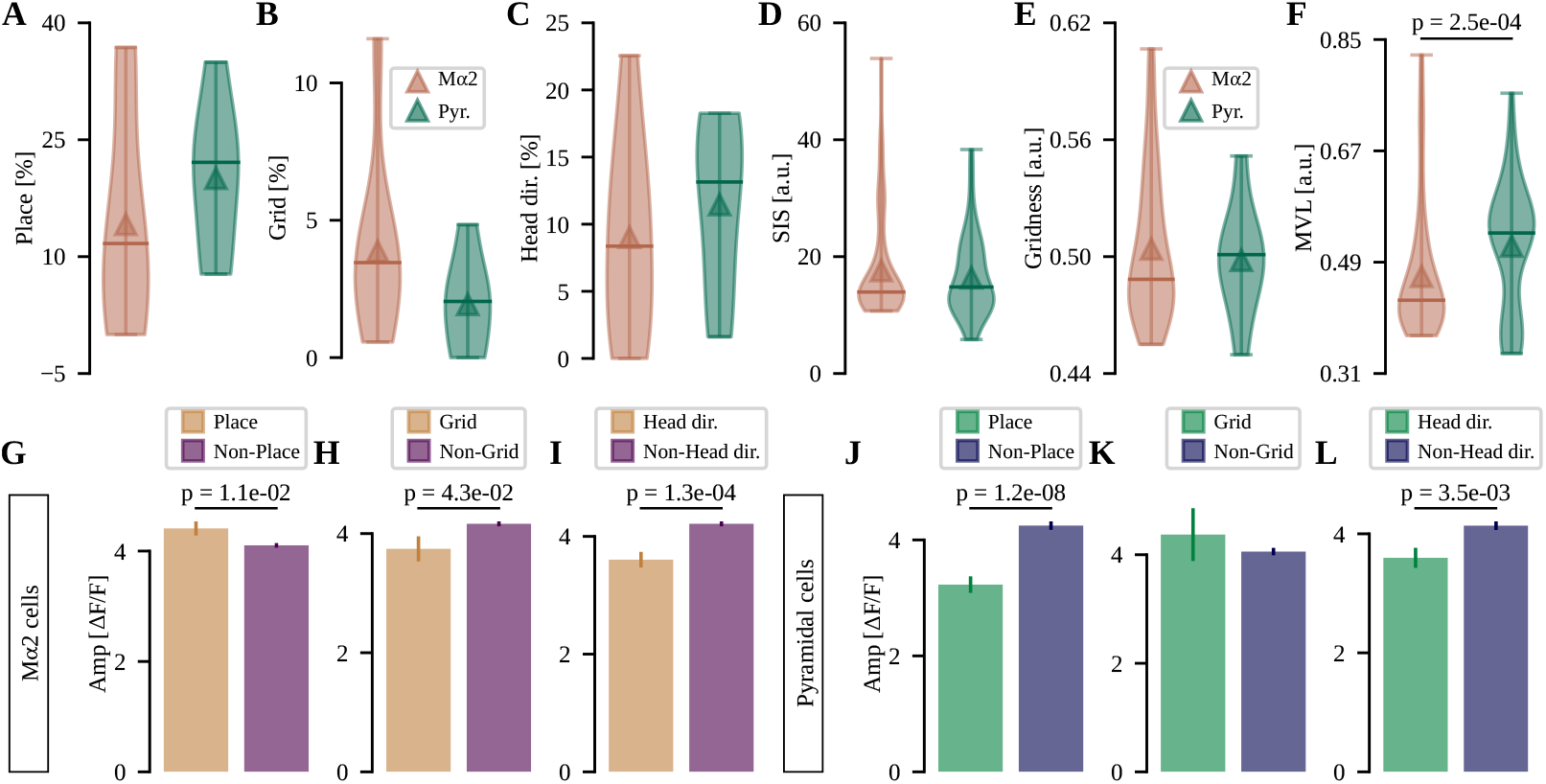
Pyramidal head direction-tuned cells are more strongly tuned than M*α*2 head direction-tuned cells. A-C) Percentage of place, grid and head-direction cells for M*α*2 (orange) and pyramidal cells (green). D-F) Comparison of SIS, gridness and MVL between M*α*2 and pyramidal cells. G-I) Amplitude of M*α*2 place-tuned, grid-tuned and head direction-tuned cells activity, represented as mean ± SEM, compared to non-place-tuned, non-grid-tuned and non-head direction-tuned cells, respectively. J-L) Same as G-I for pyramidal cells.

### Tuned cells in M*α*2 and pyramidal neurons show different activity levels from untuned cells

Finally, we examined whether activity amplitude differs between tuned and untuned cells. We averaged all cell activity during the task to assess mean activity levels across the population. Our results show that M*α*2 place-tuned cells exhibit significantly higher mean activity than non-place-tuned cells (Mann-Whitney U eff. size = 0.072; p = 1.1e-02; Figure 4G), indicating an association between place coding and elevated activity in these cells. In contrast, grid-tuned and head direction-tuned cells displayed lower mean activity than non-tuned cells (eff. size > 0.057; p < 4.3e-02; Figure 4H-I), suggesting that grid and head direction coding may be associated with reduced activity levels in M*α*2 cells. In contrast, pyramidal place-tuned cells exhibited lower mean activity than non-place-tuned cells (eff. size = 0.242; p = 1.2e-08; Figure 4J) and no difference was observed between grid-tuned and non-grid-tuned cells (Figure 4K). However, similar to M*α*2 cells, pyramidal head direction-tuned cells also displayed lower mean activity compared to non-head direction-tuned cells in pyramidal cells (eff. size = 0.124; p = 3.5e-03; Figure 4L).

## Discussion

This study offers novel insights into the neural mechanisms of spatial processing within the primary motor cortex, revealing that both layer 5 M1 M*α*2 cells and pyramidal neurons are capable of processing spatial information, including place, head direction, and grid cell representations. Our findings build upon previous research suggesting that the hippocampus plays a pivotal role in modulating neocortical activity, particularly in integrating spatial and temporal information [44, 45]. Furthermore, the hippocampal-retrosplenial cortex-M2 pathway has been implicated in facilitating spatial navigation by storing environmental information [13, 46] and previous studies have shown that M*α*2 cells in the primary motor cortex receive inputs from the M2 area [11, 47]. Whereas spinal cord central pattern generators can autonomously regulate locomotion, this process is ultimately orchestrated by integrating descending motor commands from the primary motor cortex [48], which suggest a participation of M1 during spatial navigation.

Studies have shown participation of different cortical areas in spatial navigation highlighting the importance of cortical modulation in regulating place cell activity, for example, lesions to the prefrontal cortex have been shown to alter the spatial firing properties of hippocampal place cells by reducing their mean firing rates [49]. Furthermore, activation of the prefrontal cortex enhances the firing activity of place cells within their respective fields without affecting their excitability or bursting propensity [50]. In addition to the prefrontal cortex, previous findings have also implicated the parietal cortex in spatial coding. Cells in the posterior parietal cortex exhibit directional selectivity, for example firing in response to forward motion only when moving away from the maze center, and direction-selective posterior parietal cortex cells whose firing rates are modulated by specific types of movement, such as left turns, right turns, and forward motion [51]. The converging evidence suggests that multiple hippocampal-entorhinal pathways are involved in conveying information about an animal’s current location to the posterior parietal cortex [52], which then relays this information on to the motor planning system through connections with premotor and motor regions [53].

Inhibitory interneurons play a role in shaping the activity of pyramidal cells, thereby contributing to the intricate organization of neural networks. The overrepresentation of cells that are both place-tuned and head direction-tuned relative to chance, combined with the greater amount of spatial information encoded by pyramidal cell activity compared to M*α*2 cell activity, provides further evidence that M*α*2 and pyramidal cells employ a distinct coding strategy for spatial information processing. While pyramidal neurons have been shown to exhibit place-like activity in various cortical regions, including the visual cortex [14], studies using tetrode recordings often face challenges to precisely identify the specific cell types involved, suggesting that the encoding of spatial information may extend beyond pyramidal neurons to other cell types. The interneurons activity has been previously demonstrated to encode spatial information, for example, while engaging in exploratory behavior on a linear track, GABAergic interneurons exhibit spatially selective firing and phase precession, indicating their involvement in a precisely co-ordinated local interaction with pyramidal cells [54]. Here, we show a subpopulation of Martinotti cells exhibiting spatially-tuned activity, and those cells synchronize pyramidal cell activity [26], which suggest that those cells may shape pyramidal cell encoding of spatial information. Interestingly, our findings are similar to previous observations in the hippocampus, where inhibitory interneurons exhibit spatially-tuned firing properties similar to those of excitatory neurons [23]. Our findings suggest that although interneurons can display similar spatial coding properties, such as place field size, grid size and grid angle, they might have distinct spatial coding mechanisms, as shown by the broader head direction tuning compared to pyramidal cells and the increased place-tuned activity compared to untuned cells.

Although an interesting finding, the higher mean activity of M*α*2 place-tuned cells and the decreased activity of pyramidal place-tuned cells compared to non-tuned cells during locomotion align with the expected M1 circuitry. Martinotti cells can synchronize pyramidal cell activity and connect to multiple pyramidal cells [26]. This suggests that the observed difference in activity may be due to the need for more specific neural activation patterns in pyramidal place-tuned cells, which we hypothesize can be achieved with higher levels of lateral inhibition. If this is correct, and assuming that place-tuned M*α*2 cells are connected to place-tuned pyramidal cells, these findings would support the idea that M*α*2 cells may play a role in modulating spatial representations through their connections to pyramidal cells. Another interesting finding is that both M*α*2 and pyramidal head direction-tuned neurons exhibit reduced overall activity compared to non-head direction-tuned cells which may be attributed to the high variability in activity among these neurons. Head-direction tuned neurons display elevated activity levels only within their preferred head direction range, whereas non-tuned neurons can achieve high activity levels independently of the animal’s head orientation. This variability and flexibility may contribute to a higher average activity across non-tuned neurons.

The motor cortex relies on environmental information to adjust movements, as it is essential for effective motor control [55]. Here we found a neuronal population encoding spatial information, similar to other structures such as the visual cortex (6.1% place-tuned and 11.7% head direction-tuned, Zong et al., 2022; 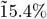 direction-specific, Vélez-Fort et al., 2018; 31-56% tuned to head-orienting movements, Guitchounts et al.,2020) and somatosensory cortex (9.6% place-tuned, 3.5% head direction-tuned and 3.6% grid-tuned, Long et al., 2001). In addition, cells in the primary somatosensory and motor cortices exhibit spatial position and orientation firing preferences in recordings from monkeys navigating a room to reach a reward spot [4], further supporting the hypothesis that spatially-tuned activity is present in the M1.

When evaluating the independence of spatial features, we found that place-tuning and head direction-tuning are not independent for both M*α*2 and pyramidal cells. These results corroborate previous findings that report neurons with both place and head direction tuning, such as those in the retrosplenial cortex, where both place cells and direction-dependent place cells have been described [12]. Similarly, in the posterior parietal cortex, neural activity of single cells can be accurately predicted based on the mouse’s position and head orientation during a virtual navigation and decision task [18]. Furthermore, lesioning the anterodorsal thalamic nuclei or the postsubiculum - regions with a large proportion of head direction cells - disrupts the stability of place cell fields in the hippocampus [56], indicating that orientation and position processing are closely linked. These findings suggest that spatial processing in the motor cortex may involve coordinated interactions between place and head direction signals, consistent with broader evidence of spatially integrated neural coding in the brain.

Although the motor cortex can store spatial information that could be important for adjusting the movements during navigation, it has a major role initiating and executing the dexterous and non-dexterous steps of skilled movement [20, 21] rather than explicit spatial mapping, resulting in a limited population of neurons directly coding for space. Our findings highlight the complex neural mechanisms underlying spatial processing in the primary motor cortex. The distinct coding strategies employed by pyramidal cells and M*α*2 interneurons underscore the importance of understanding the functional roles of different cell types in processing spatial information. Further research is necessary to elucidate the specific mechanisms underlying spatial representation, particularly in M*α*2 place-tuned and head direction-tuned cells. Moreover, the role of these cell types in other aspects of spatial cognition, such as navigation and memory formation, remains an open question. The discovery of distinct neural coding strategies employed by different cell types within the motor cortex emphasize the significance of investigating the functional roles of these cells in processing spatial information.

## Author Contributions

BC: Methodology, Data curation, Validation, Visualization, Original draft preparation, Conceptualization, Writing - Reviewing and Editing.

TM: Software, Formal analysis, Visualization, Original draft preparation, Conceptualization, Writing - Reviewing and Editing.

KK: Resources, Conceptualization, Writing - Reviewing and Editing.

## Acknowledgments

We thank Uppsala University behavioral facility (UUBF) for support

## Data Availability Statement

The datasets generated and/or analyzed in the current study are available on request.

